# How to Build a Fruit: Transcriptomics of a Novel Fruit Type in the Brassiceae

**DOI:** 10.1101/492371

**Authors:** Shane Carey, Kerrin Mendler, Jocelyn C. Hall

## Abstract

Comparative gene expression studies are invaluable for predicting how existing genetic pathways may be modified or redeployed to produce novel and variable phenotypes. Fruits are ecologically important organs because of their impact on plant fitness and seed dispersal, modifications in which results in morphological variation across species. A novel fruit type in the Brassicaceae known as heteroarthrocarpy enables distinct dispersal methods in a single fruit through segmentation via a lateral joint and variable dehiscence at maturity. Given the close relationship to Arabidopsis, species that exhibit heteroarthrocarpy are powerful models to elucidate how differences in gene expression of a fruit patterning pathway may result in novel fruit types. Transcriptomes of distal, joint, and proximal regions from *Erucaria erucarioides* and *Cakile lanceolata* were analyzed to elucidate within and between species differences in whole transcriptome, gene ontology, and fruit patterning expression profiles. Whole transcriptome expression profiles vary between fruit regions in patterns that are consistent with fruit anatomy. These transcriptomic variances do not correlate with changes in gene ontology, as they remain generally stable within and between both species. Upstream regulators in the fruit patterning pathway, *FILAMENTOUS FLOWER* and *YABBY3*, are expressed in the distal and proximal regions of *E. erucarioides*, but not in the joint, implicating alterations in the pathway in heteroarthrocarpic fruits. Downstream gene, *INDEHISCENT*, is significantly upregulated in the abscissing joint region of *C. lanceolata*, which suggests repurposing of valve margin genes for novel joint disarticulation in an otherwise indehiscent fruit. In summary, these data are consistent with modifications in fruit patterning genes producing heteroarthrocarpic fruits through different components of the pathway relative to other indehiscent, non-heteroarthrocarpic, species within the family. Our understanding of fruit development in Arabidopsis is now extended to atypical siliques within the Brassicaceae, facilitating future studies on seed shattering in important Brassicaceous crops and pernicious weeds.

## Introduction

Studying gene expression patterns across plant structures and species can elucidate how their modification may produce morphological variation (1,2). Fruits are diverse and ecologically relevant plant structures to investigate because their morphological variation determines how their seeds are dispersed (3,4). There are multitudinous fruit morphologies in nature, and they are often categorized as fleshy or dry. Fleshy fruits are distributed primarily by animals, as the seeds are discarded before or after consuming. Dry fruits however, may be dispersed by animals, wind, or water. Dry fruits are further classified by whether they are dehiscent, releasing seeds into the environment, or indehiscent, releasing seeds in a protected fruit wall propagule. Thus, variation in fruit morphology is directly tied to differences in dispersal capabilities.

*Arabidopsis thaliana* (Brassicaceae) is the premier model for dry dehiscent fruits. Arabidopsis fruits have been characterized from gynoecium formation to seed release, and many genes responsible for fruit development are described, as are their interactions (5–7). This knowledge forms a basis of comparison in the investigation of complex trait morphologies that diverge from Arabidopsis, especially amongst close relatives e.g., the loss of dehiscence in many species across the Brassicaceae (1).

Brassicaceae fruits vary markedly in shape, structure, and size (1,8). Their variation in dehiscence is a focal point for research because it fundamentally changes fruit structure, subsequently affecting dispersal and diversification (9). A prerequisite for exploring how differences in fruit morphology are achieved across the Brassicaceae is familiarity with both the fruit structure and underlying genetic pathways in *Arabidopsis* (10,11). *Arabidopsis* fruits, hereafter referred to as typical siliques, are composed of five basic elements: valve, replum, seeds, septum, and valve margins. The valve, synonymous with ovary wall in Arabidopsis, is the outermost tissue of the fruit that protects the developing seeds and is separated from the replum at maturity to release seeds. The replum is the persistent placental tissue to which the seeds are attached. The septum, which connects to the replum, divides the fruit into two locules or chambers. The valve and replum are separated by the valve margin, which consists of a lignification and separation layer. Thus, proper fruit formation relies on the establishment of medial (replum) and lateral (valves and valve margin) components (12). As the fruit dries, tension is created via the lignified layer, which facilitates the separation of the valves from the replum at the separation layer (13). This general morphology is stable across most dehiscent members of Brassicaceae (1).

The causal factors for dehiscence have been well characterized in Arabidopsis (14–17), with proper formation and positioning of the valve margin being a key to this process. The valve margin pathway is essential for spatial regulation and development of valve, replum, and valve margin tissues (11,18–23). Briefly, *FRUITFULL* (*FUL*) and *REPLUMLESS* (*RPL*), as well as other upstream regulators, restrict the expression of the valve margin genes to two cell layers between the valve and replum, respectively. The valve margin genes, *SHATTERPROOF 1/2* (*SHP1/2*), *INDEHISCENT*(*IND*), *SPATULA* (*SPT*), and *ALCATRAZ* (*ALC*), are responsible for the formation of the valve margin, specifically of the separation and lignification layers that control dehiscence (Fig 1). Upstream regulators of *FUL* and *RPL*, e.g., *APETALA2* (*AP2*), *FILAMENTOUS FLOWER* (*FIL*), *YABBY3* (*YAB3*), and *JAGGED* (*JAG*) are also key to precise positioning of the valve margin because they tightly regulate downstream processes. In sum, replum and valve genes function in an antagonistic manner to ensure proper formation of these regions of the fruit(12).

**Figure 1.**
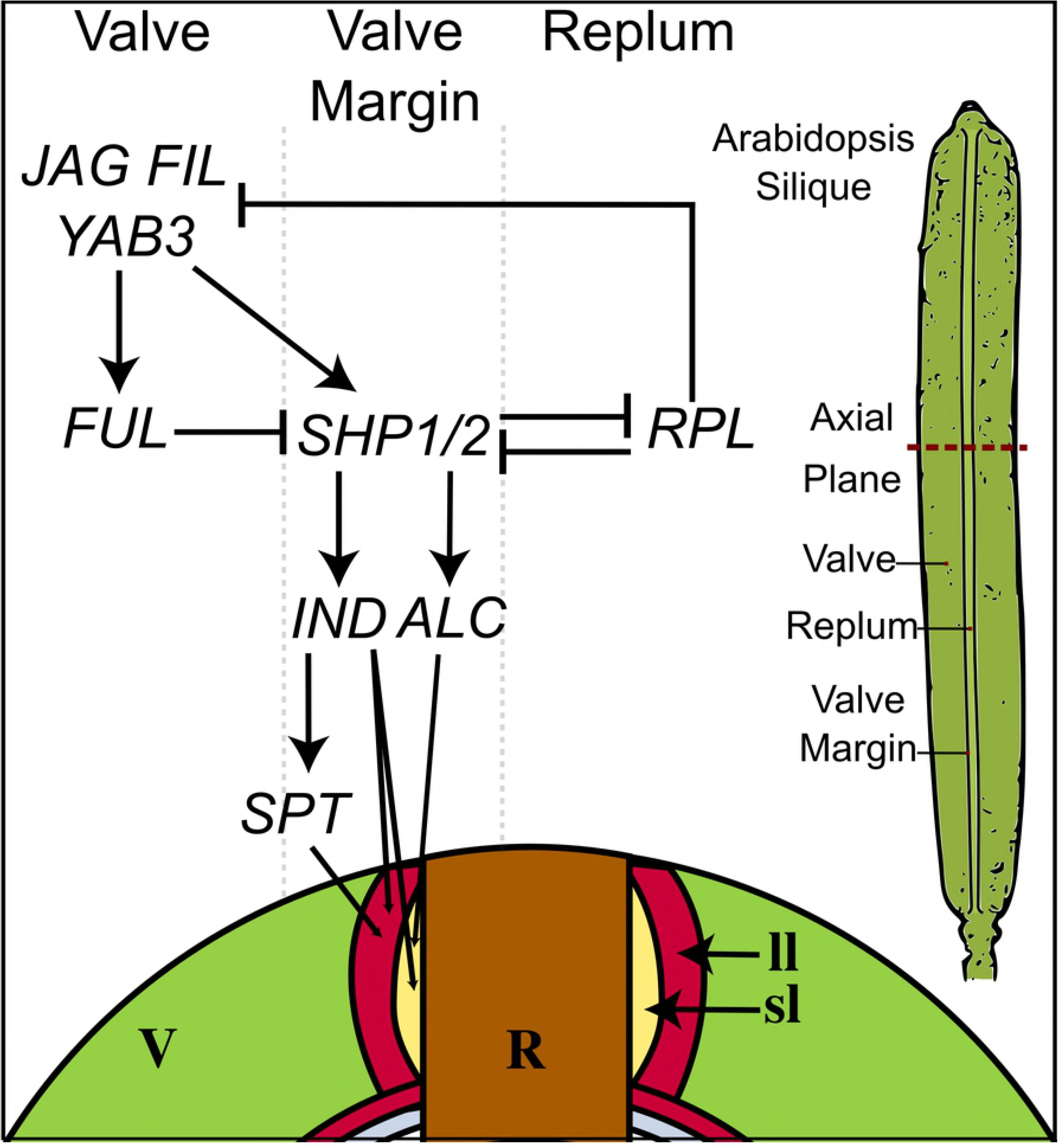
Diagram of simplified valve margin pathway for fruit dehiscence in *Arabidopsis thaliana*; valve margin. R, replum. Sl, separation layer. ll, lignification layer. Valve margin = sl + ll. Modified from data available in (10-11,14) and figure 2 (36).

Most of the *Arabidopsis* valve margin genes are pleiotropic and many of them share indehiscence as a phenotype of mutation. For example, a mutation in any of the following genes results in indehiscent fruits in *Arabidopsis: SHP1/2, SPT, ALC* and *IND* (24–27). Overexpression of *FUL* or *NO TRANSMITTING TRACT* (*NTT*) also results in indehiscent fruits (28,29); *FUL* overexpression completely suppresses *SHP1/2*, resulting in reduced lignification in the en*b* layer and reduced valve margin formation; overexpression of NTT phenocopies the *ful* mutation resulting in valve margin specific genes being expressed throughout valve. In summary, a modification of many components in this pathway results in a loss of dehiscence. Because indehiscence is observed in at least 20 different lineages across the family, it is likely that this phenotype evolved via multiple modifications to this pathway (30). As such, there is no singular alteration to the fruit patterning pathway implicated in this shift for all tribes.

To date, little is known about the genetic basis of indehiscence in the Brassicaceae, although it is currently being bridged by studies in taxa with varying indehiscent morphologies. Recently, a study demonstrated a deviation in expression of eight key genes between pod shatter sensitive species and shatter resistant species of *Brassica* and *Sinapis* (2). In *Lepidium*, there has been an evolutionary shift from dehiscence to indehiscence, e.g., valve margin genes that are conserved between the dehiscent *L. campestre* and *Arabidopsis* have been lost in the indehiscent *L. apellianum* (31,32). Upregulation in upstream regulator *AP2* has been suggested as a factor in this indehiscence (32).

A notable morphological adaptation is the evolution of a complex fruit type known as heteroarthrocarpy, which is only found in some members of the tribe Brassiceae (30,33,34). This modified silique is defined by the presence of a variably abscising central joint, an indehiscent distal region, and a variably dehiscent proximal region (Fig 2). As such, this novel morphology offers an opportunity to investigate fruit variation beyond shifts from dehiscent to indehiscent. Anatomically, heteroarthrocarpic fruits appear most like *Arabidopsis* siliques in their proximal regions, varying by a lack of a valve margin cell layer in indehiscent variants (35–37). There are three described variations of heteroarthrocarpy: a non-abscising joint with a dehiscent proximal region, an abscising joint with an indehiscent proximal region or an abscising joint with a dehiscent proximal region (36). These subtypes have evolved multiple times, perhaps as a bet hedging strategy in response to selective pressure from hostile desert environments (9,37). Heteroarthrocarpic subtypes may be developmental enablers that have facilitated changes in fruit morphology across the tribe, which would explain heteroarthrocarpy’s evolutionary lability (36). Regardless of lability, all types are linked by the mechanism in which seeds from the same fruit are released by different means. In other words, the joint is the novel and unifying feature of heteroarthrocarpy (36).

**Figure 2.**
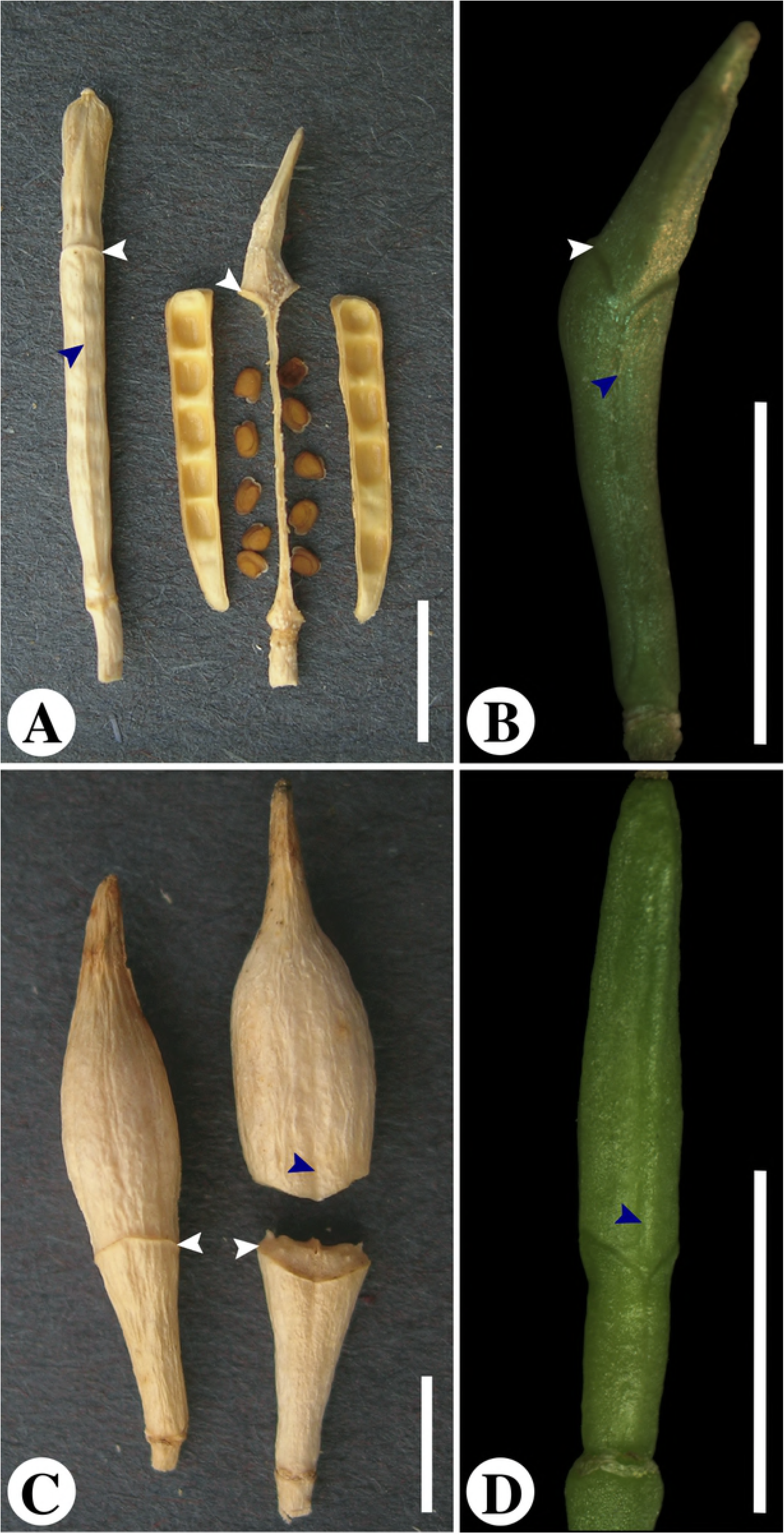
Mature and young heteroarthrocarpic fruits. (A), Mature *Erucaria erucarioides* fruit in lateral view before dehiscence (left), and medial view after dehiscence (right). (B), Young *E. erucarioides* fruit in medial view. C, *Cakile lanceolata* fruit in lateral view before dehiscence (left), and medial view after joint abscission (right). (D), Young *C. lanceolata* fruit in medial view; Modified from figure 1 (36). White arrows indicate joint region; blue arrows indicate replum. Scale bars = 5mm

A comparison of expression patterns between heteroarthrocarpic subtypes is potentially informative for formulating hypotheses about its evolutionary origins. *Erucaria erucarioides* and *Cakile lanceolata*, hereafter referred to as *Erucaria* and *Cakile*, are two well-studied representatives for heteroarthrocarpy because of their close relation and divergent subtypes (Fig 2) (9,10,36,37). In previous studies it was hypothesized that the formation of heteroarthrocarpy is the result of repositioning of the valve margin, such that the valve is only present in the proximal region of the fruit, unlike in Arabidopsis where it is found in the entire ovary (36). In other words, the joint is the distal portion of the valve margin. This hypothesis was partially supported by comparative gene expression data of some, but not all, genes in the valve margin pathway using a candidate gene approach (10). However, that study did not definitively determine how the pathway has been repositioned because it did not investigate upstream genes. Candidate gene approaches will, by design, overlook non-targeted genes, and a lack of in situ hybridization does not necessarily indicate a lack of expression. Further, the basis of the joint remains unknown.

No study to date has investigated transcriptional variation of heteroarthrocarpic fruits sectioned transversely into distal, joint and proximal regions. This approach is complementary to prior research because it quantifies expression of all transcripts in discrete regions of a whole system. Expression profiles from these regions will elucidate broad patterns and potentially identify key players involved in the formation of heteroarthrocarpy. They will clarify unique and shared gene expression patterns between and within *Erucaria* and *Cakile*, and will set the groundwork for future research regarding the evolution of the joint. Herein, the objective is to uncover transcript patterns, unique or shared, between and within, two variant heteroarthrocarpic species. We expect gene expression to be consistent with anatomical features within fruits, and that expression of fruit patterning transcripts will be consistent with repositioning of the valve margin in heteroarthrocarpy.

## Materials and Methods

### Plant material

Seeds from *Erucaria erucarioides* (Coss. and Durieu) Müll.Berol and *Cakile lanceolata* (Willd.) O.E.Schulz were obtained from the late César Gómez-Campo’s and KEW royal botanical garden’s seed collections, respectively. Vouchers for *Cakile* and *Erucaria* have been deposited in the Vascular Plant Herbarium at the University of Alberta, and the Harvard University Herbaria, respectively. Seeds were germinated in 1% agar and transferred to clay pots containing a 2:1 soil (Sungro sunshine mix #4, Agawam, MA, USA) to perlite mixture. Plants were grown under a 16/8-hour light/dark schedule at 24°C with scheduled watering in the University of Alberta, Department of Biological Sciences, growth chambers.

Distal, joint, and proximal regions from 10mm fruits (~10 days post fertilization) were collected and flash frozen in liquid nitrogen prior to storage at -80°C. Distal and proximal regions were classified as all tissue ~1mm above or below the joint, and the joint is remaining tissue between distal and proximal regions (Fig 2). The 10mm fruit size is roughly equivalent to Arabidopsis stage 17A fruits (7), which go through elongation and cell expansion before maturity. This size was chosen to capture late stage valve margin gene expression because the valve margin is easily distinguished at this stage, and an increase in lignification is observed in key layers, e.g., en*b*. (36).

### RNA isolation and cDNA library preparation

RNA was extracted from frozen tissue using manual grinding and a Qiagen RNeasy micro kit (Hilden, Germany) with the following amendments to protocol: RNA was incubated in nuclease free water for five minutes prior to elution, and this eluate was spun through the same extraction column to maximize RNA yield. RNA concentration was verified using a Nanodrop ND-1000 spectrophotometer (Software version 3.1.2), and quality was confirmed using the Agilent 2100 bioanalyzer (Software version B.02.09.SI720). All cDNA samples were set at the same concentration of the most dilute RNA extraction. Samples were processed using the Illumina TruSeq stranded mRNA LT sample prep kit RS-122-2101 (California, U.S.), and the procedure was followed as described in the low sample protocol. The mRNA from each sample was isolated and purified using AMPure XP magnetic beads (Agencourt; Beverly, Massachusetts) before primary and secondary strand cDNA synthesis. Unique Illumina adapters were ligated, and each sample was PCR amplified before validation. Samples were normalized, pooled, and sequenced by the center for applied genetics (TCAG) facilities of the Toronto Sick Kids hospital, Ontario, Canada.

### De novo transcript assembly, differential expression, and annotation

Raw reads were trimmed and quality checked using Trim Galore! (Version 0.4.1) (38) and FastQC (Version 0.11.3) (39) then assembled using Trinity (Version 2.2.0) (40). Corset (Version 1.0.6) (41) was used to estimate contig abundance by grouping contigs into representative gene clusters as the first step of the differential expression analysis. Contigs are defined as continuous overlapping paired-end reads. Next, edgeR (Version 3.6.2) (42,43) was used to perform pairwise differential expression analysis of Trinity gene, Trinity contig, and Corset clusters between proximal, joint, and distal regions of the same species. Genes, contigs, and clusters were classified as significantly differentially expressed if log2(fold-change)> 2 and the False Discovery Rate (FDR)-corrected p-value (α) < 0.05. The analyze_diff_expr.pl script, provided with Trinity, was used to generate z-score heatmaps of all significantly differentially expressed contig clustered transcripts (α) < 0.05. A z-score is used to indicate how many standard deviations a value is above the mean. The transcriptomes were annotated using the Basic Local Alignment Search Tool (BLAST) (44) algorithm on a local copy of both the National Center for Biotechnology Information (NCBI) non-redundant protein (nr) database and The Arabidopsis Information Resource (TAIR) database (45). BLASTx (E-value<10^-10^) was used to identify highly similar sequences, and transcripts with the highest bit-score from the TAIR database were used as representative transcripts for heatmap generation. Whole transcriptome and fruit patterning heatmaps were generated using ggplot2 (46) and ggplot in R, respectively (Version 3.4.2) (47).

### Orthologous Clustering

Orthofinder (Version 1.1.8) (48) was used to match orthologous transcripts from unfiltered *Erucaria* and *Cakile* transcriptomes. Orthogroups containing transcripts from both species as well as top BLAST matches for fruit patterning genes of interest were used to generate heatmaps. For Venn diagram generation, high-throughput sequencing (HTS) (49) filtered transcripts, sorted by regions, were translated to longest open reading frame (ORF) protein fasta files using TransDecoder (Version 5.0.0) (50). These files were uploaded for comparison using the Orthovenn webserver (51). HTS filtering was used to reduce file size due to the web server upload limit, and to reduce the number of insubstantial transcripts.

### Gene ontology

Transcriptome fasta files from *Erucaria* and *Cakile* were imported to BLAST2GO (Version 2.8) (52). Annotation files were exported and filtered to generate gene ontology (GO) terms for each region and species. These GO terms were used to produce graphs containing transcriptome hits for chosen terms. Terms were chosen based on searches for lignin, abscission, dehiscence, specific hormone keywords, and top hits. For comparison between transcriptomes, the log2 of selected GO term counts were divided over the log2 of all GO term counts (log2(n)/log2(N)).

## Results

### De novo Assembly of *Erucaria* and *Cakile* Transcriptome Data

RNA-seq libraries were constructed from 9 total replicates of triplicate distal, proximal, and joint regions. RNA samples from segmented fruits of two distinct plants were combined before sequencing to achieve optimal yield for library preparation. Sequencing from both libraries averaged 27.41 and 29.41 million paired-reads for *Erucaria* and *Cakile*, respectively. After quality trimming read counts were reduced to 27.36 million and 28.36 million high quality reads, respectively. Inter-quartile ranges per base were minimally 33 for *Erucaria* for the first 5 base pairs, and minimally 32 in the 90^th^ percentile; *Cakile’s* inter-quartile ranges were minimally 33 for the first 5 base pairs, and minimally 29 in the 90^th^ percentile.

The transcriptome from *Erucaria* had an average contig length of 942.83, and *Cakile’s* had an average length of 877.15. The total transcript count for *Erucaria* and Cakile was 227,530 and 314,194 reads, respectively (Table 1). Corset cluster counts averaged 365,257 (*Erucaria*) and 436,177 (*Cakile*). Notably, the first replicate for *Cakile* had a read count of 269,732, which is minimally 130,000 fewer than replicate 2 and 3. This inconsistency may have caused some issues in downstream analyses, but overall, both transcriptomes were of adequate quality and read-depth.

**Table 1.** Statistics for de novo Trinity assembly of *Erucaria erucarioides* and *Cakile lanceolata* pairwise reads.

| All (Longest Isoform) | *Erucaria* | *Cakile* |
| --- | --- | --- |
| **N50** | 1544 (1017) | 1464 (835) |
| **Median Contig Length** | 578 (374) | 517 (330) |
| **Average Contig length** | 942.83 (656.94) | 877.15 (577.55) |
| **Total Assembled bases** | 214,521,562 (92,098,767) | 275,595,508 (108,815,069) |
| **Total Trinity Genes** | 140194 | 184945 |
| **Total Trinity Transcripts** | 227530 | 314194 |
| **GC%** | 41.89 | 42.05 |

### Annotation of Assembled Transcripts

Both transcriptomes were compared to the nr and TAIR peptide database using a BLASTx algorithm, and all downstream analyses used the TAIR10 annotation for facilitated comparison to Arabidopsis. A total of 254,592 (*Cakile*) and 213,757 (*Erucaria*) transcripts with e-values≤10^-5^ were matched to the TAIR10 database with multiple transcripts matches per gene. The GO analysis averaged 8,644 and 8,941 terms for *Erucaria* and *Cakile*, respectively. The top 15 GO terms consisted of 11 cellular component, three molecular function, and one biological process. Nucleus, plasma membrane, and protein binding were the top three terms, all of which are biological processes (Fig S1).

The majority of selected orthogroups were similar between and within species (lignin, abscission, and dehiscence processes, and hormone response) (Fig 3). Exceptions include: cell wall modification related to abscission, general abscission, and catabolic lignification. *Cakile* has a greater ratio of cell wall modification processes and a lower ratio of general abscission processes relative to *Erucaria. Erucaria* has a higher ratio of catabolic lignification processes in the joint region despite having similar ratios relative to *Cakile* in the distal and proximal regions (Fig 3). Overall, the GO analysis results are consistent between and within species.

**Figure 3.**
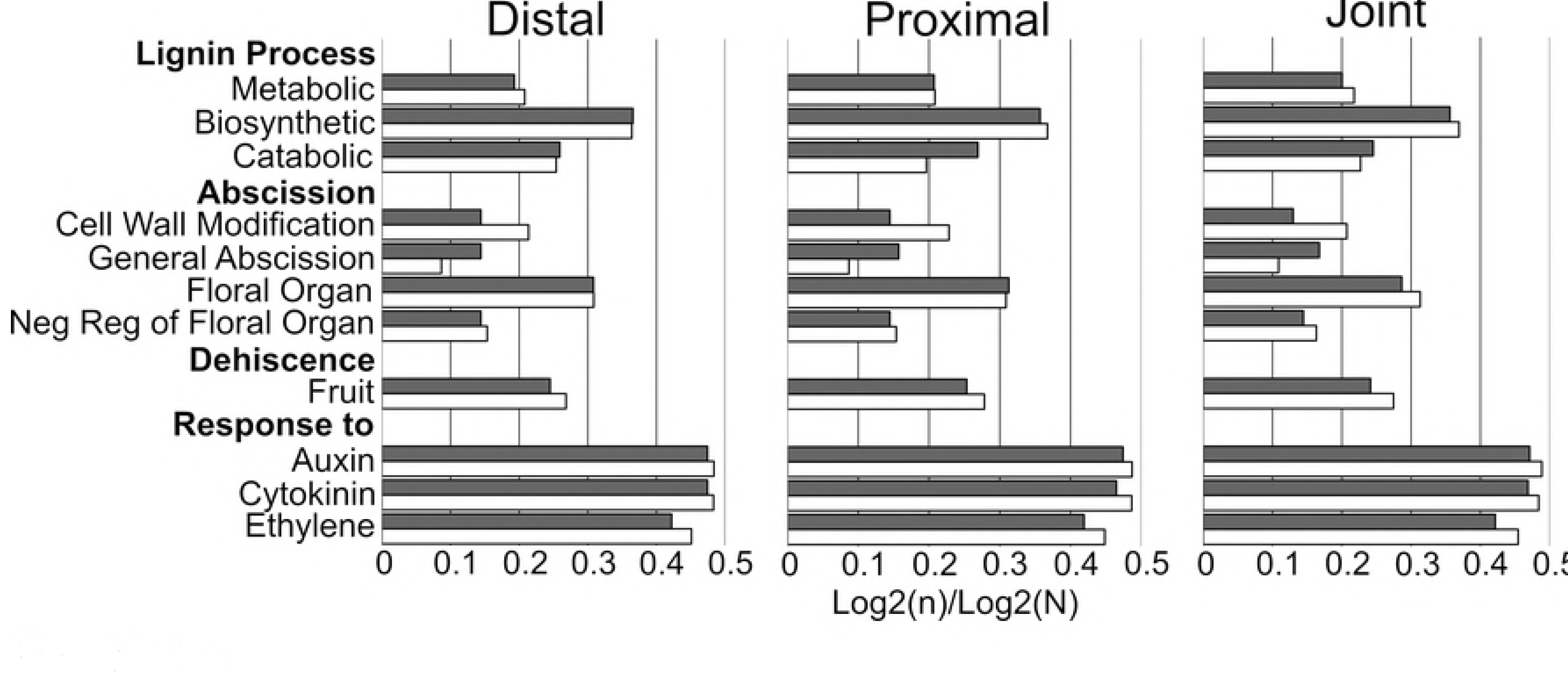
Graph of select Gene Ontology (GO) terms for *Erucaria erucarioides* and *Cakile lanceolata*. Sample(n) and total(N) raw counts log2 transformed for interspecies comparison. GO terms chosen based on search terms: lignin, abscission, dehiscence, and response to hormone.

Additional results from OrthoVenn showed minimal difference in orthologous clustering within species, but some differences between species (Fig 4). There are a greater number of shared clusters between the proximal and distal regions in *Erucaria* (2548) than *Cakile* (2306) despite *Cakile* having substantially more overall clusters than *Erucaria* (50,003 vs 32,757). Additionally, there are fewer clusters unique to the joint for *Cakile* (21) than *Erucaria* (112). In sum, there are fewer orthologous clusters in common within regions of *Cakile* fruits than within regions of *Erucaria* fruits.

**Figure 4.**
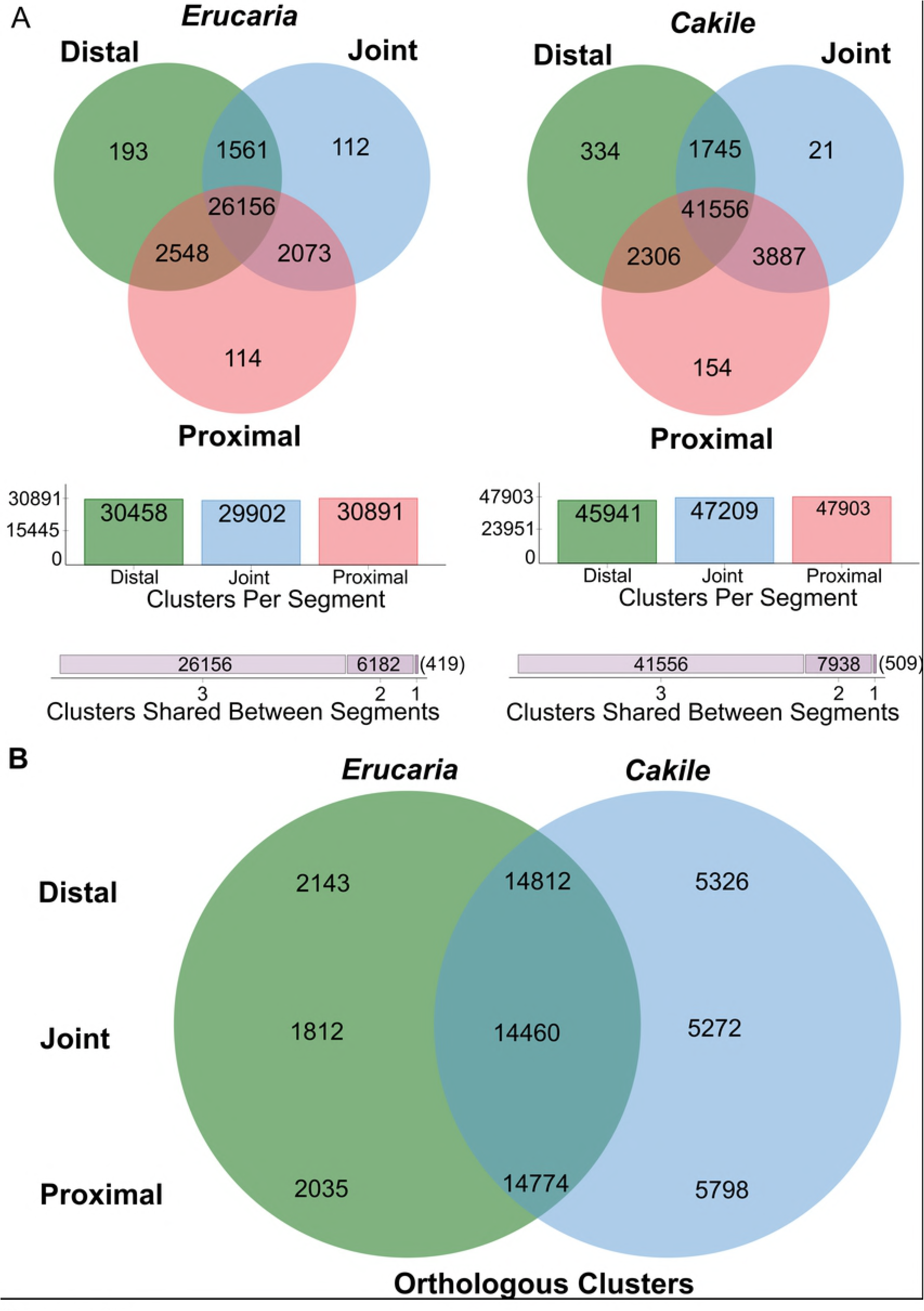
Venn diagrams of three-way and pairwise High Throughput Sequencing (HTS) filtered transcripts for *Erucaria erucarioides* and *Cakile lanceolata* transcriptomes. (A), Three-way Venn diagrams of *Erucaria* and *Cakile* orthologous clusters for distal, joint, and proximal regions. (B), Pairwise Venn diagrams of *Erucaria* and *Cakile* orthologue-clustered transcripts (*Erucaria* region vs *Cakile* region).

*Cakile* shares a greater number of representative transcripts from the valve margin pathway with *Erucaria* (12) than *Erucaria* does with *Cakile* (7), i.e., more representative BLAST transcripts from *Cakile* are orthologous with transcripts from the *Erucaria* transcriptome than vice versa. Of the representative valve margin gene transcripts, only two are orthologous between both species, *ASYMETRIC LEAVES 2* (*AS2*) and *SHP2*. (Fig 5 and 6). Representative transcripts are those with the highest bit-score after a BLAST search against the TAIR database.

**Figure 5.**
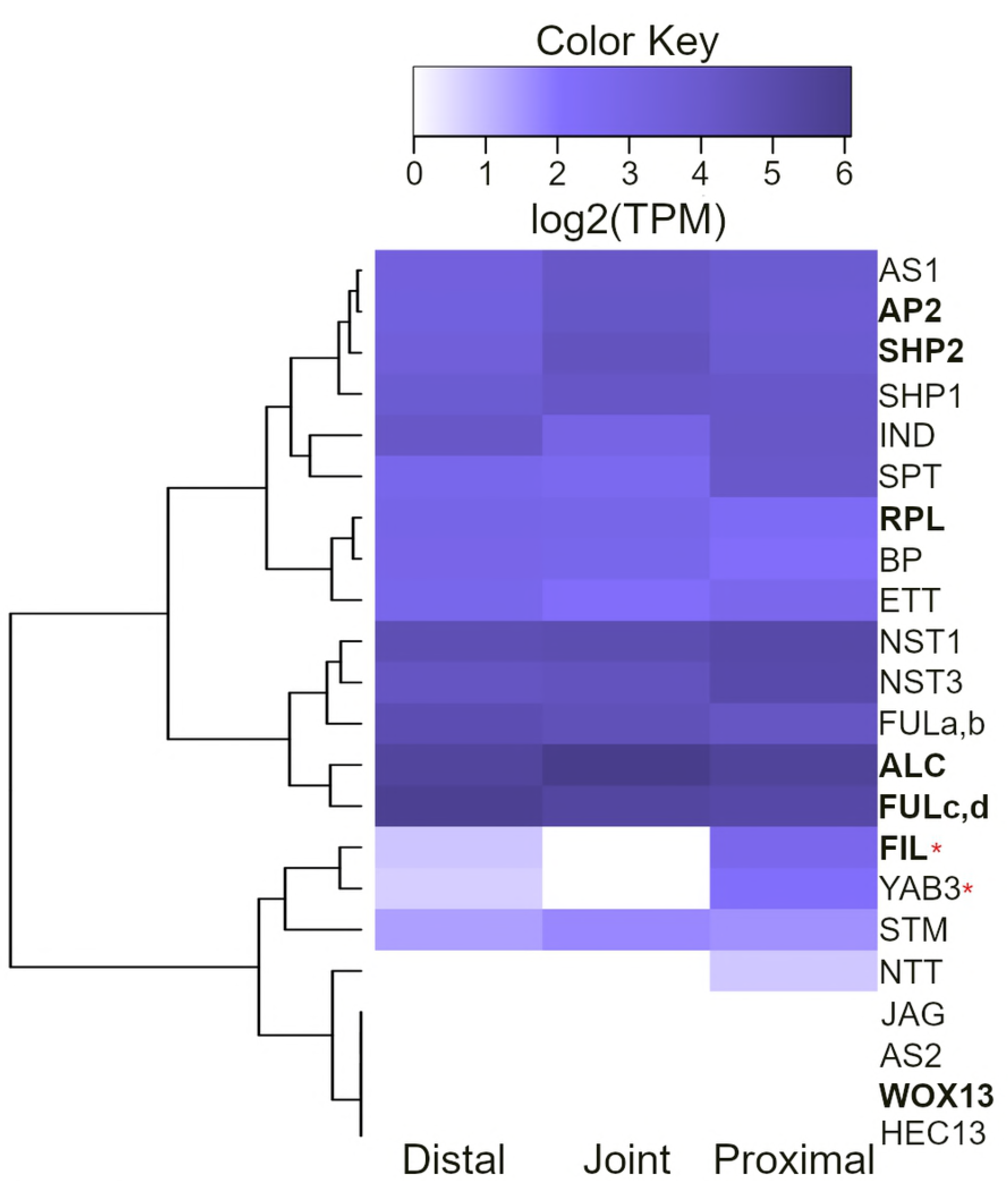
Heatmap of edgeR contig clustered transcripts from *Erucaria erucarioides* expressed in log2 TPM with TMM normalization. Representative transcripts based off largest bitscore hit against TAIR database. Bolding indicates shared orthogroup with *Cakile lanceolata*. TPM, Transcripts Per Million; TMM, Trimmed Mean of M-values. *FULa,b,c,d* are copies of *FUL* that are present in some species across the Brassicaceae (72). Asterisks indicate significant differential expression between proximal and joint region (FDR-corrected α=0.01)

**Figure 6.**
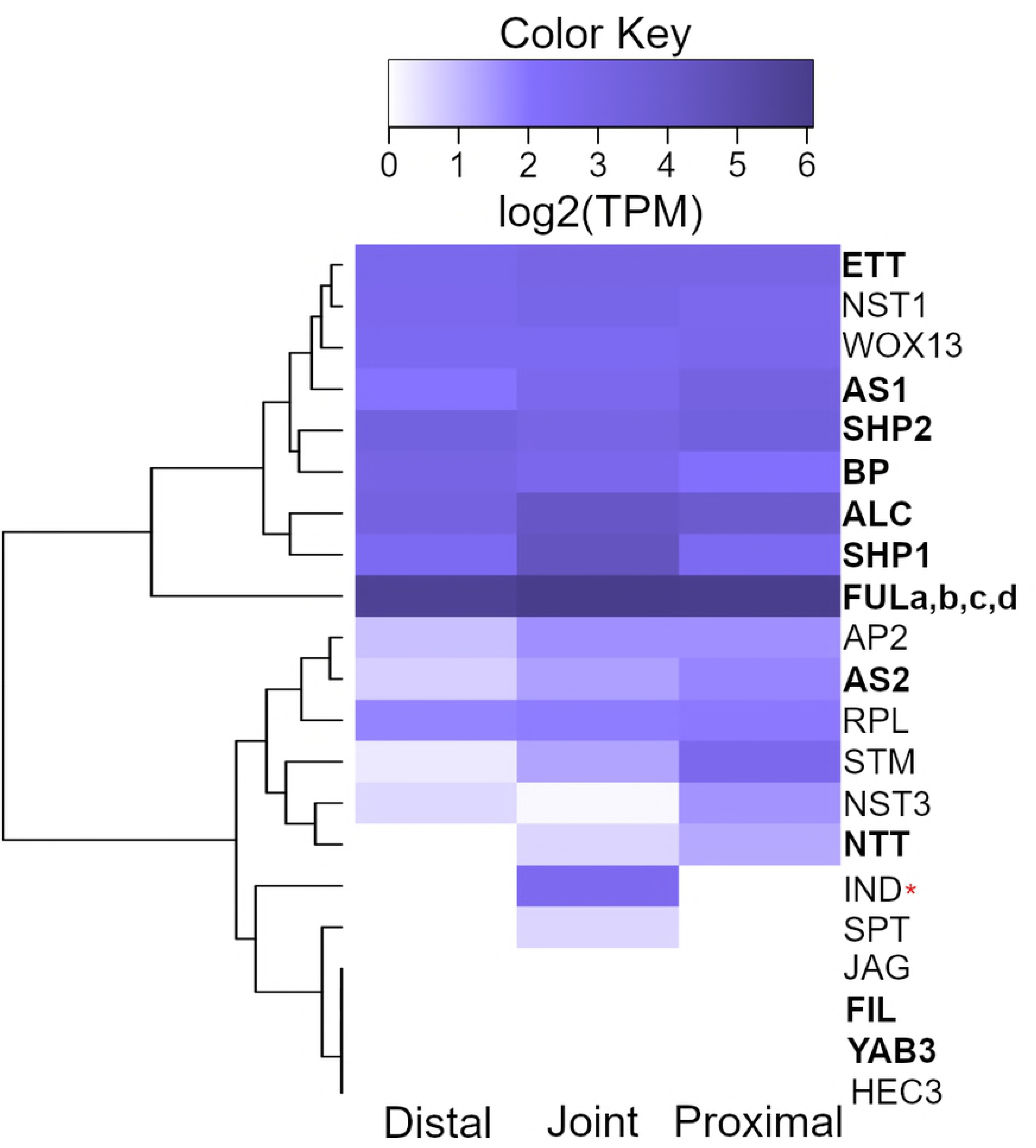
Heatmap of edgeR contig clustered transcripts from *Cakile lanceolata* expressed in log2 TPM with TMM normalization. Representative transcripts based off largest bitscore hit against TAIR database. Bolding indicates shared orthogroup with *Cakile lanceolata*. TPM, Transcripts Per Million; TMM, Trimmed Mean of M-values. *FULa,b,c,d* are copies of *FUL* that are present in some species across the Brassicaceae (72). Asterisks indicate significant differential expression between distal and joint region (FDR-corrected α=0.05)

### Identification of Differentially Expressed Transcripts in 10mm fruit

For whole transcriptome comparison, two heatmaps of significant pairwise differentially expressed transcripts (α = 0.01) were generated (Figs 7 and 8). Contig clustering was chosen for this analysis because it is a more conservative estimation of significant differential expression at the transcript level, i.e., there are a greater number of transcripts being compared with more stringent FDR correction relative to corset clustering. Values were then converted to z-score to facilitate interspecies comparison, and for visual clarity. The joint and proximal regions of *Erucaria* are most alike in expression and are both dissimilar to the distal region (Fig 7). All three regions in *Cakile* have different expression patterns, and the distal region has a relatively large inter-replicate variance (Fig 8). There are 15,345 (*Erucaria*) and 74 (*Cakile*) significantly differentially expressed (SDE) transcripts in each transcriptome. There were no SDE *Cakile* transcripts with FDR-adjusted p-values < 0.01. The low number of SDE genes between *Cakile* regions indicates a lack of regional distinction in terms of transcript expression. These data demonstrate a large difference in significant differential expression between the distal region relative to the joint and proximal region in *Erucaria*, and little significant variation between all three *Cakile* regions.

**Figure 7.**
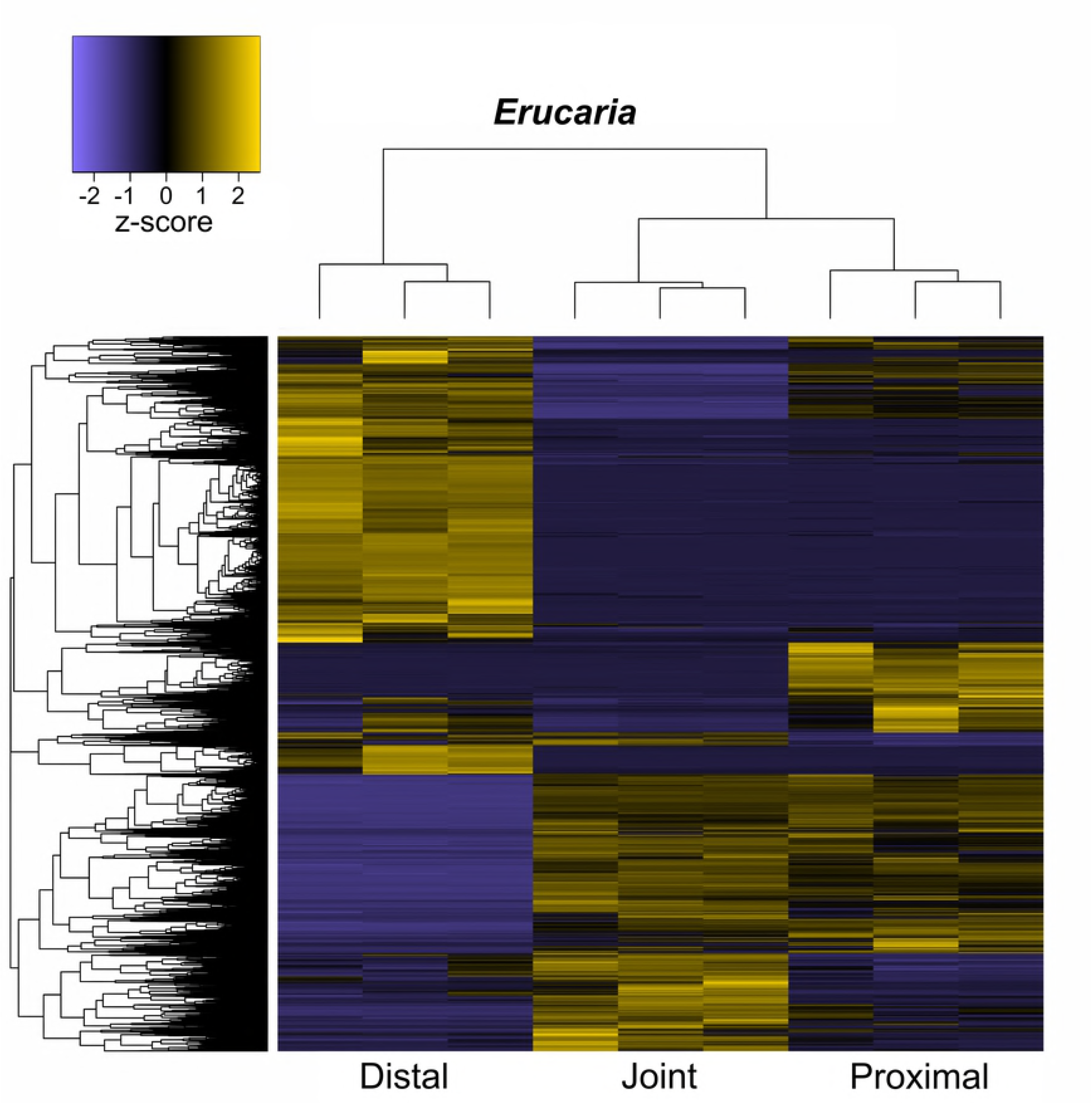
Heatmap of all significant edgeR contig clustered transcripts in *Erucaria erucarioides,* expressed as z-scores (FDR-corrected α=0.01). Row and column dendrograms indicate clustering of transcripts (n=15,345) and biological replicates (n=3 per region), respectively.

**Figure 8.**
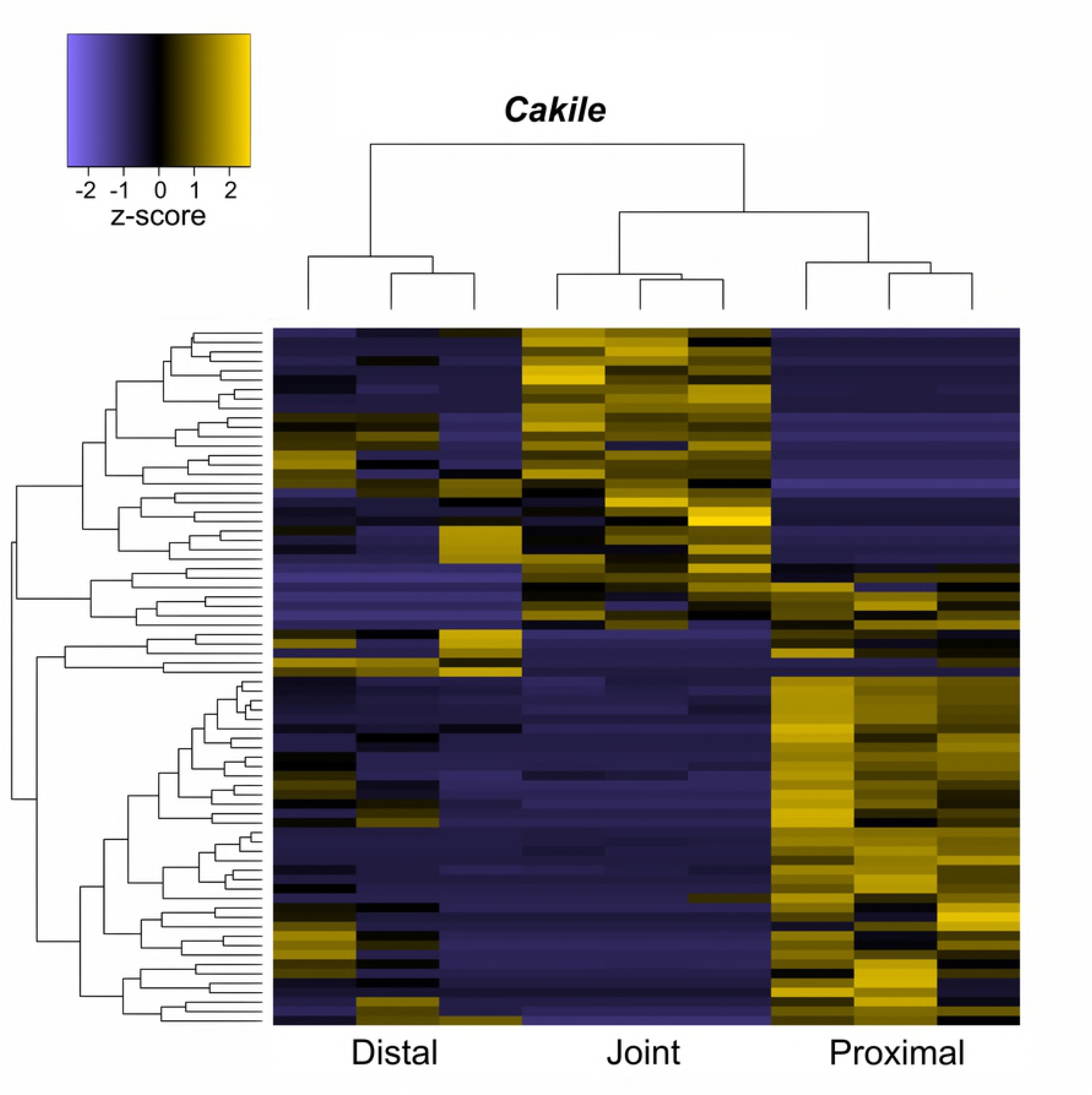
Heatmap of all significant edgeR contig clustered transcripts in *Cakile lanceolata*, expressed as z-scores (FDR-corrected α=0.01). Row and column dendrograms indicate clustering of transcripts (n=74) and biological replicates (n=3 per region), respectively.

We compared expression profiles of 21 genes important for valve margin formation and positioning in Arabidopsis (2,11,16,53–64) (Fig 5 and 6). Contig clustered transcripts were also chosen for this analysis based on matches against the TAIR database. Most fruit patterning genes for both species have no significant differences in expression across all regions, except for *FIL* and *YAB3* which were significantly upregulated in the distal region relative to the joint in *Erucaria*, and *IND* which is significantly upregulated in the joint relative to both the distal and proximal regions in *Cakile*. Upstream regulators *FIL* and *YAB3* are not expressed in late stage *Cakile* fruits, despite global expression in *Erucaria* fruits. Downstream regulator *IND* is expressed in the whole fruit in *Erucaria*, but only in the joint region of *Cakile* (Fig 5 and 6).

## Discussion

### Gene ontology of heteroarthrocarpic fruits

Overall, GO terms within fruits and between species are similar (Fig 3 and S1), as expected, because all sections and replicates are from developing fruit with shared components (e.g., ovary wall, septum). Because dehiscence is susceptible to misexpression and loss of function mutations in the valve margin pathway (21,24–28), broad changes in gene ontology are unnecessary to explain heteroarthrocarpy. Additionally, GO analyses of top terms do not usually vary between closely related species (65,66). However, similarities in gene ontology do not imply similarity between all expressed transcripts, so variation of just a few transcripts may still be the driving factor behind heteroarthrocarpy.

### Global transcript expression of heteroarthrocarpic fruits are consistent with anatomy

Transcript expression patterns are consistent with anatomical variances within and between fruits. The distal region of *Erucaria* has opposing transcript expression relative to both its joint and proximal regions (Fig 7), i.e., when transcripts are upregulated distally in *Erucaria* they are downregulated proximally. This pattern is consistent with heteroarthrocarpic fruit anatomy, as distal regions contain no valve or valve margin, and proximal regions have both (36). In contrast, all regions of *Cakile* have variable transcript expression, with the clearest distinction between the proximal and joint regions, i.e., when genes are upregulated proximally they will be downregulated in the joint (Fig 8). As with *Erucaria*, expression profiles in *Cakile* vary in a manner consistent with anatomy. Superficially, one would expect the *Cakile* silique to have similar expression between all regions because the entire fruit is indehiscent, however the distal region of *Cakile* is more like the distal region of *Erucaria* than to its own proximal region (36). Further, its abscising joint is anatomically reminiscent to a valve margin (36). Abscission zones are also found between septum and seeds, and they too share similar anatomy and expression to typical silique valve margins (67). Heteroarthrocarpic distal regions are unlike indehiscent non-heteroarthrocarpic siliques such as *L. appelianum*, because heteroarthrocarpic distal regions have no remnant valve margin in contrast to indehiscence observed in *Lepidium* and the proximal region of *Cakile* (32,36). Thus, we expect different expression patterns within heteroarthrocarpic fruits, as well as between heteroarthrocarpic and non-heteroarthrocarpic fruits. In summary, there is a clear difference between distal and proximal expression profiles for both *Erucaria* and *Cakile*, which is consistent with a repositioning of the valve margin, i.e., the distal region is quite distinct from the proximal region due to the lack of valve margin, or its remnant, in the distal region. This consistency is further explored by analysis of fruit patterning transcript expression involved in valve margin formation.

### Fruit patterning genes

Despite the substantial differences in anatomy, most valve margin genes reveal similar expression patterns across fruits in both *Erucaria* and *Cakile* (Fig 5 and 6). These differences are initially surprising because previous studies showed variation in expression patterns across fruits with in-situ hybridization (10). *EeFUL1*, one of two *FUL* homologs found in *Erucaria*, was previously shown to only be expressed in the proximal region in earlier stages of carpel development (10), but all *FUL* transcripts are expressed across all regions in this study of later stage development (Fig 5). This discrepancy may be due to dynamic gene expression at different stages or because our methodology cannot distinguish within region differences (e.g., genes expressed in valve but not replum), so differences within regions cannot be distinguished. In contrast to *EeFUL1*, our data are consistent with a previous publication which demonstrated that other fruit patterning genes have broader expression domains than found in Arabidopsis (10). *EeALC* and *EeIND* and *ClALC* were expressed in the septum of *Erucaria* and *Cakile,* respectively, which is found throughout all regions sampled in this study. The replum is also found throughout all sampled regions of *Erucaria* and *Cakile*, so expression patterns of pleiotropic genes, e.g., *AP2*, show broader expression patterns than expected in valve and valve margin alone.

It is a compelling finding that upstream regulators *FIL/YAB3* and *JAG* have variable expression across *Erucaria* (Fig 5). These three genes positively regulate expression of *FUL* and valve margin genes in Arabidopsis such that their cooperative function has been designated together as *JAG/FIL* activity (12). Our data suggest a decoupling of this cooperation in heteroarthrocarpic fruits because these three genes do not exhibit the same expression patterns across *Erucaria* fruits (Fig 5 and 6). That is, no expression of *JAG* was detected in any region of either species at this stage. *FIL* and *YAB3* showed differently expression patterns across fruits of *Erucaria*, but neither were detected in *Cakile*. It is important to note that the double mutant of *fil/yab3* in Arabidopsis have fruits that are remarkably reminiscent of heteroarthrocarpy: they lack valve margin in the distal region of fruit while maintaining ovary wall identity (11). In contrast to heteroarthrocarpy, these mutants have ectopic valve margin in the proximal region of their fruits (11). As these genes exhibit different patterns across *Cakile* and *Erucaria* and are expressed in both proximal and distal regions of *Erucaria*, heteroarthrocarpy cannot be explained by a simple lack of expression of these key regulators. Further, *FIL/YAB* are absent in the joint region of *Erucaria* (Fig 5), which is confounding since the joint contains small portions of both proximal and distal regions, an unavoidable consequence of segmentation during tissue collection. Nonetheless, deviation in expression patterns of these upstream regulators between Arabidopsis and heteroarthrocarpic fruits implicates variation in their expression profiles in the origin of heteroarthrocarpy.

When exploring heteroarthrocarpy, we need to consider fruit patterning beyond the basal-apical differences that distinguish distal, joint, and proximal regions. That is, the lateral (valve and valve margin) and medial (replum) patterning is maintained in heteroarthrocarpic fruits whereas the apical-basal is not. In other words, not only is replum tissue present in distal, joint, and proximal regions of heteroarthrocarpic fruits, it is appropriately sized. *FIL/YAB3* and *JAG* function antagonistically with replum promoting gene, *WUSCHEL RELATED HOMEOBOX 13* (*WOX13*), which positively regulates *RPL* in turn. This interaction is necessary for proper medial-lateral formation of Arabidopsis fruits. Further, *ASYMMETRIC LEAVES1* (*AS1*) and *AS2* collaborate with *JAG/FIL* function as promoters of lateral factors (12). The loss of both *AS1*/2 and *JAG/FIL* in Arabidopsis results in dramatic medial-lateral differences and substantially enlarged repla, which is interestingly more pronounced in the basal portion of the fruit (12,68). As *AS1*/2 and *AS1* are expressed throughout *Cakile* and *Erucaria* regions, respectively, this pattern suggests that *AS1* alone is sufficient for proper replum (aka medial-lateral) formation in heteroarthrocarpic fruits. In other words, the collaboration between *JAG/FIL* function and *AS1*/2 is not maintained in heteroarthrocarpic fruits. Further, in at least *Cakile JAG/FIL* activity is non-detectable in the entire fruit, at least at later stages of development. Thus, it appears that some redundancy in lateral-medial patterning of Arabidopsis fruits has been lost in heteroarthrocarpic fruits, while simultaneously gaining apical-basal differences.

### Valve margin pathway recruitment and abscission in the Cakile joint

The fruit of *Cakile* is distinct in that the joint abscises (disarticulates) at maturity. The joint, which represents the distal portion of the valve margin, thus represents a novel abscission zone in *Cakile*, completely separating the distal portion of the fruit. This is an unusual feature of certain heteroarthrocarpic subtypes, as there is no equivalent abscission zone in Arabidopsis. Our data strongly implicate the recruitment of downstream valve margin genes as responsible for joint abscission, although how that zone is positioned remains elusive. *IND* is significantly upregulated in joint region (Fig 6) and is primarily responsible for formation of separation and lignification layers in typical siliques (24,26), a juxtaposition of cell types also observed in the abscising joint region. Its presence in the joint may be due to a co-option of downstream valve margin pathway genes to facilitate formation of the joint abscission zone. Similar co-option is observed in seed abscission zones, although these zones typically involve *SEEDSTICK* (*STK*) in lieu of *SHP*, and the functionally similar transcription factor *HEC3* in lieu of *IND* (67). *SHP1/2* and *ALC* expression are both consistent with this co-option, as they are expressed in all three regions (Fig 6). Additionally, *SPT* expression is consistent with expression of *IND*, as expected from its downstream role in valve margin formation. (Fig 6) (14). Further, both representative transcripts are among the 21 unique orthologous clusters in the joint of *Cakile* (Fig 4). This pattern is consistent with in situ hybridization data that showed *SHP2* expressed in septum and ovules of *Cakile*, and in ovules of *Erucaria* (10). Thus, the likely function of *SHP1/2* and *ALC* in the joint region would be to promote expression of *IND* (*SHP1/2*), and the formation of the separation layer (*ALC*). What is unusual about joint abscission is that for the joint to separate, the distal and proximal regions of the replum must also separate. This expression pattern then implies that the mechanism used to physically separate valve from replum may also be in play for replum in the joint region. Taken together with anatomical studies, our data strongly suggests that there is a repurposing of the valve margin pathway in an otherwise indehiscent *Cakile* fruit, and that this pathway may be capable of initializing disarticulation in multiple tissue types.

## Conclusion

Transcriptomic expression from late stage *Erucaria* and *Cakile* fruits is consistent with some conservation and some deviation of the valve margin pathway, specifically in upstream regulation, e.g., *YAB/FIL/JAG*. Thus, different upstream regulators are implicated in the loss of dehiscence in Brassiceae relative to *Lepidium*, where *AP2* is likely responsible (32). Loss of expression of *YAB/FIL/JAG* in Arabidopsis results in differing apical and basal phenotypes, which may help to explain the apical/basal differences in heteroarthrocarpic fruits (11). Further, heteroarthrocarpic fruits likely recruit the same mechanism used in valve and seed abscission for joint abscission (Fig 6). Functional tests are necessary to confirm whether redeployment of *FIL/YAB3, IND*, and possibly *SPT* have key roles in the origin of heteroarthrocarpy as well as joint abscission.

There have been multiple whole genome duplications in the Brassicales, which has resulted in many polyploids within the Brassicaceae family (69–71). We considered the possibility of transcriptional differences between gene copies in distal, joint, and proximal regions that were undetected because we were unable to determine copy number in our transcriptome. For example, there are four copies of *FUL* in the Brassiceae (72), but each potential *FUL* copy had multiple hits from the same transcripts in both transcriptomes, so there is no definitive answer about copy number and expression (Fig 5 and 6). That is, we could not confirm or refute subfunctionalization of some fruit patterning genes as having a role in the origin of heteroarthrocarpy. An analysis of multiple transcripts for every fruit patterning gene showed generally similar expression for each, but further analyses are needed to determine if neo/subfunctionalization plays a role in heteroarthrocarpy.

Understanding the nature of heteroarthrocarpy, and how it relates to fruit development in Arabidopsis, will facilitate future studies on seed shattering in important Brassicaceous crops, and pernicious heteroarthrocarpic weeds. Further, these studies inform on the origin of important variation in seed packaging and dispersal capabilities.

## Acknowledgements

The authors thank Navjot Singh and Erin Yue for their invaluable help in fruit collection. We also thank NSERC for providing the funding required to complete this research.

## Author Contributions Statements

SC and JH contributed concept and project design. SC and KM contributed to plant care, RNA extraction and cDNA library preparation. KM designed scripts and was lead in initial bioinformatic analyses; SC completed later analyses using scripts produced by KM. SC wrote the first manuscript draft; SC and JH wrote subsequent manuscript drafts. All authors contributed to revision and proofreading of the final submitted version.

## Conflict of interest statement

There are no conflicts of interest to report.

---

**Figure S1.**
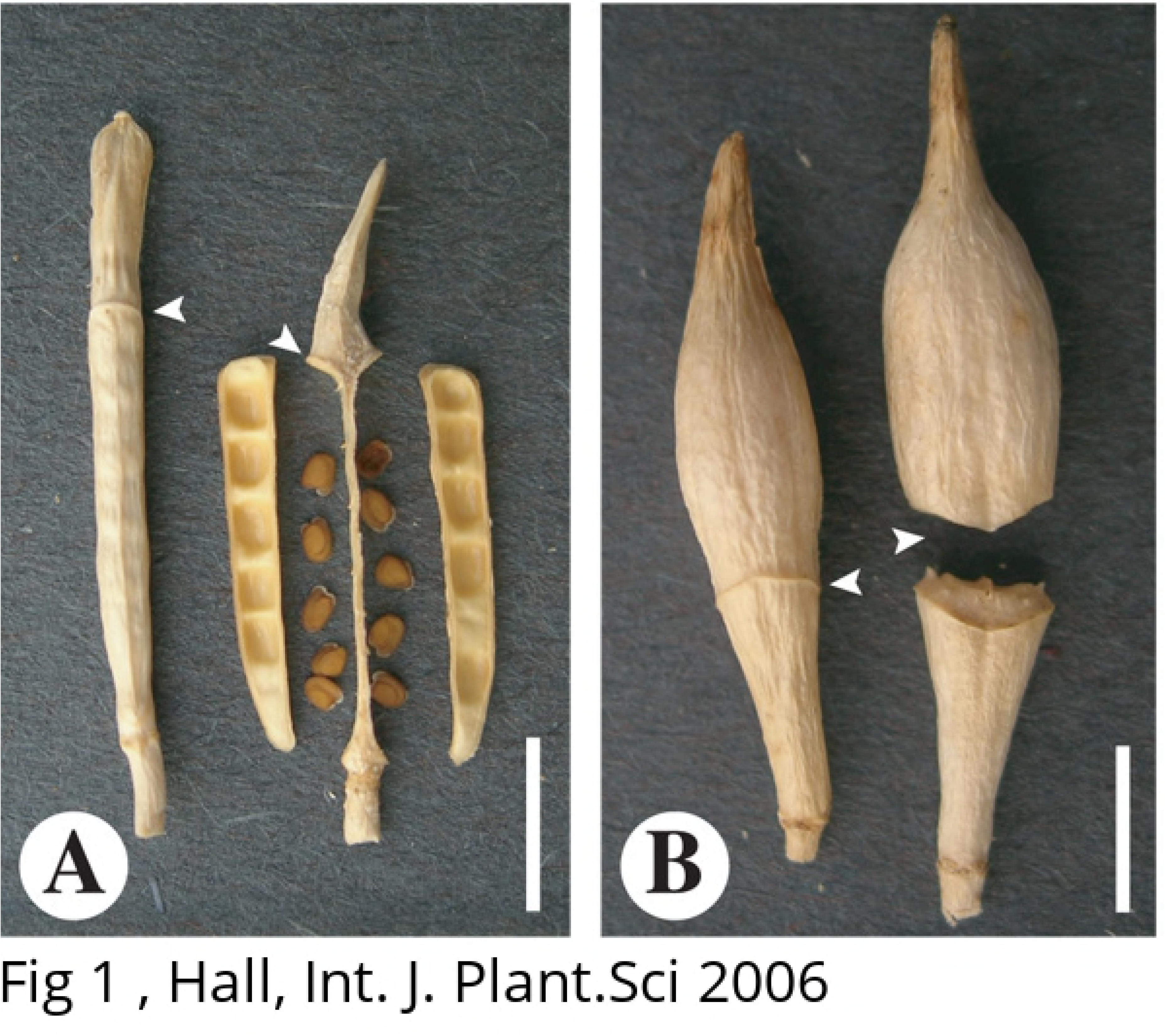
Graph of top Gene Ontology (GO) terms for *Erucaria erucarioides* and *Cakile lanceolata*. Sample(n) and total(N) raw counts were log2 transformed for interspecies comparison.

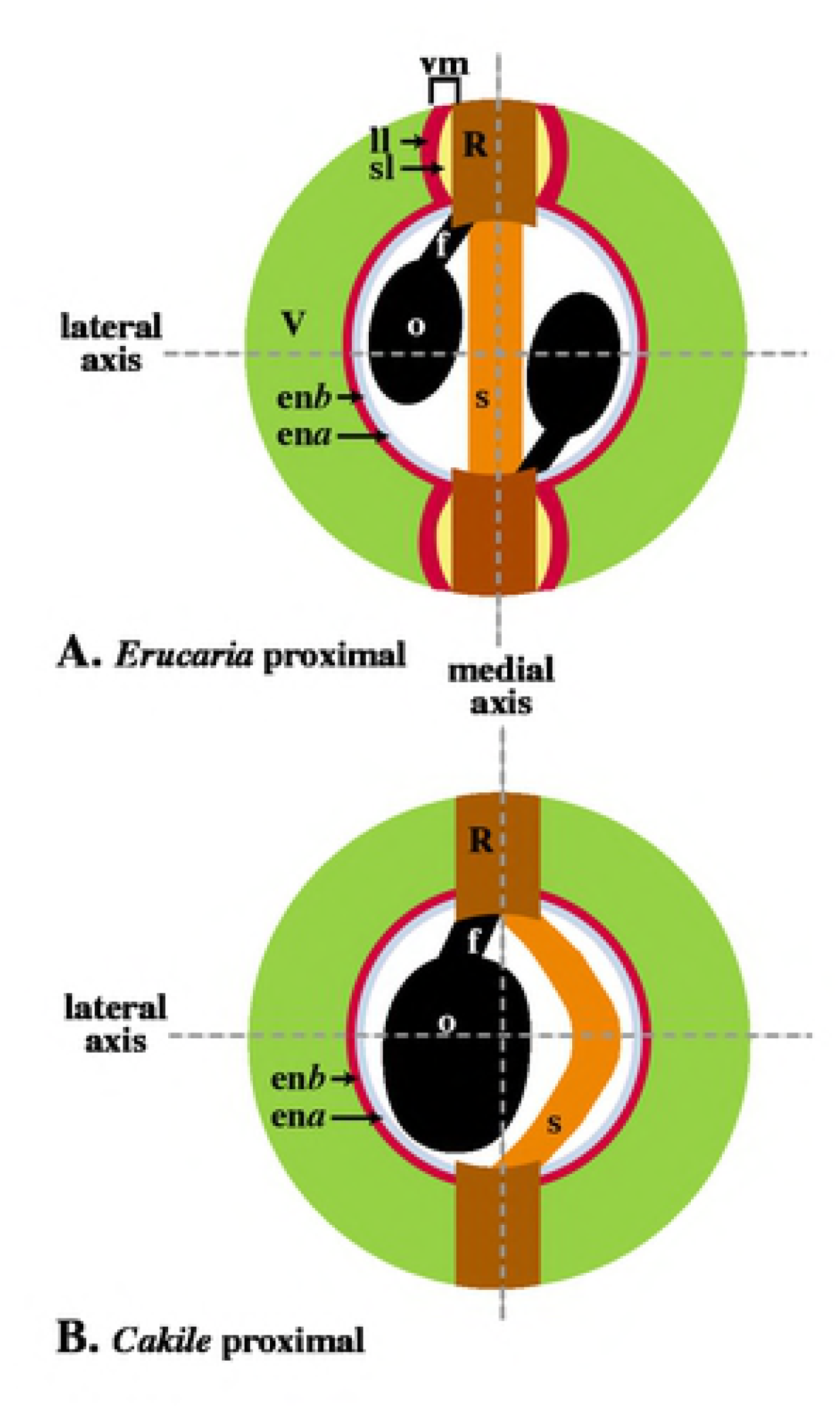

## References

1. Łangowski Ł, Stacey N, Østergaard L. Diversification of fruit shape in the Brassicaceae family. Plant Reprod. 2016;29:149–63.

2. Zhang Y, Shen YY, Wu XM, Wang JB. The basis of pod dehiscence: anatomical traits of the dehiscence zone and expression of eight pod shatter-related genes in four species of Brassicaceae. Biol Plant. 2016;60:343–54.

3. Gautier-hion AA, Duplantier J, Quris R, Feer F, Sourd C, Decoux J, et al. Fruit Characters as a Basis of Fruit Choice and Seed Dispersal in a Tropical Forest Vertebrate Community. Oecologia. 1985;65:324–37.

4. Janson CH. Adaptation of Fruit Morphology to Dispersal Agents in a Neotropical Forest Published by: American Association for the Advancement of Science Stable URL: http://www.jstor.org/stable/1690124 Adaptation of Fruit Morphology to Dispersal Agents in a Neotropi. Adv Sci. 1983;219:187–9.

5. Ferrándiz C, Pelaez S, Yanofsky MF. C Ontrol of C Arpel and F Ruit D Evelopment. Annu Rev Biochem. 1999;321–54.

6. Robles P, Pelaz S. Flower and fruit development in *Arabidopsis thaliana*. Int J Dev Biol. 2005;49:633–43.

7. Roeder A, Yanofsky M. Fruit Development in Arabidopsis. In: The Arabidopsis Book. 2006. p. 1–50.

8. Eldridge T, Ł Ł, Stacey N, Jantzen F, Moubayidin L, Sicard A, et al. Fruit shape diversity in the Brassicaceae is generated by varying patterns of anisotropy. Development. 2016; 143:3394–3406. doi: 10.1242/dev.135327.

9. Willis CG, Hall JC, Rubio De Casas R, Wang TY, Donohue K. Diversification and the evolution of dispersal ability in the tribe Brassiceae (Brassicaceae). Ann Bot. 2014;114:1675–86.

10. Avino M, Kramer EM, Donohue K, Hammel AJ, Hall JC. Understanding the basis of a novel fruit type in Brassicaceae: conservation and deviation in expression patterns of six genes. Evodevo. 2012;3: 20.

11. Dinneny JR, Weigel D, Yanofsky MF. A genetic framework for fruit patterning in *Arabidopsis thaliana*. Development. 2005;132:4687–96.

12. González-Reig S, Ripoll JJ, Vera A, Yanofsky MF, Martínez-Laborda A. Antagonistic Gene Activities Determine the Formation of Pattern Elements along the Mediolateral Axis of the Arabidopsis Fruit. PLoS Genet. 2012;8:e1003020.

13. Spence J, Vercher Y, Gates P, Harris N. “Pod shatter” in Arabidopsis thaliana, Brassica napus and B. juncea. 1996. doi: 10.1046/j.1365-2818.1996.111391.x.

14. Girin T, Paicu T, Stephenson P, Fuentes S, Korner E, O’Brien M, et al. INDEHISCENT and SPATULA Interact to Specify Carpel and Valve Margin Tissue and Thus Promote Seed Dispersal in Arabidopsis. Plant Cell. 2011;23:3641–53.

15. Girin T, Stephenson P, Goldsack CMP, Kempin SA, Perez A, Pires N, et al. Brassicaceae INDEHISCENT genes specify valve margin cell fate and repress replum formation. Plant J. 2010;63:329–38.

16. Gu Q, Ferrandiz C, Yanofsky MF, Martienssen R. The FRUITFULL MADS-box gene mediates cell differentiation during Arabidopsis fruit development. Development. 1998;125:1509–17.

17. Roeder A, Ferrándiz C, Mf Y. The Role of the REPLUMLESS Homeodomain Protein in Patterning the Arabidopsis Fruit. Cell Press. 1998;13:1630–5.

18. Ballester P, Ferrándiz C. Shattering fruits: variations on a dehiscent theme. Current Opinion in Plant Biology. 2017;35:68–75. doi: 10.1016/j.pbi.2016.11.008.

19. Chávez Montes RA, Herrera-Ubaldo H, Serwatowska J, de Folter S. Towards a comprehensive and dynamic gynoecium gene regulatory network. Curr Plant Biol. 2015. doi: 10.1016/j.cpb.2015.08.002.

20. Dinneny JR, Yanofsky MF. Drawing lines and borders: How the dehiscent fruit of Arabidopsis is patterned. BioEssays. 2005;27:42–9.

21. Ferrandiz C. Regulation of fruit dehiscence in Arabidopsis. J Exp Bot. 2002;53:2031–8.

22. Ferrándiz C, Fourquin C. Role of the FUL-SHP network in the evolution of fruit morphology and function. Journal of Experimental Botany. 2014 Aug;65:4505–13. doi: 10.1093/jxb/ert479.

23. Xu L, Palatnik J, Springer P, Mao L, Hepworth SR, Ca S, et al. Beyond the Divide: Boundaries for Patterning and Stem Cell Regulation in Plants. Front Plant Sci. 2015;6:1052. doi: 10.3389/fpls.2015.01052.

24. Groszmann M, Paicu T, Alvarez JP, Swain SM, Smyth DR. SPATULA and ALCATRAZ, are partially redundant, functionally diverging bHLH genes required for Arabidopsis gynoecium and fruit development. Plant J. 2011;68:816–29.

25. Liljegren SJ, Ditta GS, Eshed Y, Savidge B, Bowman JL, Yanofsky MF. SHATTERPROOF MADS-box genes control seed dispersal in Arabidopsis. Nature. 2000;404:766–70.

26. Liljegren SJ, Roeder AHK, Kempin SA, Gremski K, Østergaard L, Guimil S, et al. Control of fruit patterning in Arabidopsis by INDEHISCENT. Cell. 2004 Mar 19;116:843–53.

27. Kramer E, Irish V. Evolution of genetic mechanisms controlling petal development investigated the genetic mechanisms controlling petal development. 1999;336:144–8.

28. Chung KS, Lee JH, Lee JS, Ahn JH. Fruit indehiscence caused by enhanced expression of NO TRANSMITTING TRACT in *Arabidopsis thaliana*. Molecules and Cells. 2013;35:519–25. doi: 10.1007/s10059-013-0030-0.

29. Ferrándiz C, Gu Q, Martienssen R, Yanofsky MF. Redundant regulation of meristem identity and plant architecture by FRUITFULL, APETALA1 and CAULIFLOWER. Development. 2000;127:725–34.

30. Appel O, Al-Shehbaz IA. Cruciferae. In K. kubitzki and C. Bayer [eds.], The families and genera of vascular plants. In: Springer-Verlag, Berlin, Germany. 2003. p. 75–174.

31. Lenser T, Theißen G. Conservation of fruit dehiscence pathways between Lepidium campestre and *Arabidopsis thaliana* sheds light on the regulation of INDEHISCENT. Plant J. 2013;76:545–56. doi: 10.1111/tpj.12321.

32. Mühlhausen A, Lenser T, Mummenhoff K, Theißen G. Evidence that an evolutionary transition from dehiscent to indehiscent fruits in Lepidium (Brassicaceae) was caused by a change in the control of valve margin identity genes. Plant J. 2013;73:824–35. doi: 10.1111/tpj.12079.

33. Gómez-Campo C. Seedless and seeded beaks in the tribe Brassiceae. Eucarpia Crucif. 1999;21:11–2.

34. Warwick SI, Sauder CA. Phylogeny of tribe Brassiceae (Brassicaceae) based on chloroplast restriction site polymorphisms and nuclear ribosomal internal transcribed spacer and chloroplast *trn* L intron sequences. Can J Bot. 2005;83:467–83.

35. Gómez-Campo C. Morphology and morpho-taxonomy of the tribe Brassiceae, in Brassica crops and wild allies. Japan Scientific Societies press, Japan. 1980;3–31.

36. Hall JC, Tisdale TE, Donohue K, Kramer EM. Developmental Basis Of An Anatomical Novelty: Heteroarthrocarpy In Cakile Lanceolata And Erucaria Erucarioides (Brassicaceae). 2006;167:771–789

37. Hall JC, Tisdale TE, Donohue K, Wheeler A, Al-Yahya MA, Kramer EM. Convergent evolution of a complex fruit structure in the tribe Brassiceae (Brassicaceae). Am J Bot. 2011;98:1989–2003.

38. F K. Trim Galore! [Internet]. 2012. Available from: http://www.bioinformatics.babraham.ac.uk/projects/trim_galore/

39. S A. FastQC: a quality control tool for high throughput sequence data [Internet]. 2010. Available from: http://www.bioinformatics.babraham.ac.uk/projects/fastqc

40. Grabherr MG, Haas BJ, Yassour M, Levin JZ, Thompson DA, Amit I, et al. Trinity: reconstructing a full-length transcriptome without a genome from RNA-Seq data HHS Public Access. Nat Biotechnol Nat Biotechnol. 29:644–52.

41. Davidson NM, Oshlack A. Corset: Enabling differential gene expression analysis for de novo assembled transcriptomes. Genome Biol. 2014;15:1–14.

42. Robinson M, McCarthy D, Smyth G. edgeR: a Bioconductor package for differential expression analysis of digital gene expression data. Bioinformatics. 2009;26:139–40.

43. McCarthy D. Chen Y, Smyth K. Differential expression analysis of multifactor RNA-Seq experiments with respect to biological variation. Nucleic Acids Res. 40:4288–97.

44. Altschul S, Gish W, Miller W, Myers E, Lipman D. Basic local alignment search tool. J Mol Biol. 1990;215:403–10.

45. Garcia-Hernandez M, Berardini T, Chen G, Crist D, Doyle A. TAIR: a resource for intergrated Arabidopsis data. Funct Integr Genomics. 2002;2:239–53.

46. Wickham H. Ggplot2 [Internet]. 2009. Available from: http://had.co.nz/ggplot2/book%0Ahttp://link.springer.com/10.1007/978-0-387-98141-3

47. R Core Team. R: A language and environment for statistical computing. [Internet]. R Foundation for statistical Computing, Vienna, Austria. 2013. Available from: http://www.r-project.org/

48. Emms DM, Kelly S. OrthoFinder: solving fundamental biases in whole genome comparisons dramatically improves orthogroup inference accuracy. Genome Biol. 2015;16:157. doi: 10.1186/s13059-015-0721-2.

49. Rau A, Gallopin M, Celeux G, Jaffrézic F. Data-based filtering for replicated high-throughput transcriptome sequencing experiments. Bioinformatics. 2013;29:2146–52.

50. Haas BJ, Papanicolaou A, Yassour M, Grabherr M, Blood PD, Bowden J, et al. De novo transcript sequence reconstruction from RNA-seq using the Trinity platform for reference generation and analysis. Nat Protoc. 2013;8:1494.

51. Wang Y, Coleman-Derr D, Chen G, Gu YQ. OrthoVenn: A web server for genome wide comparison and annotation of orthologous clusters across multiple species. Nucleic Acids Res. 2015;43:W78–84.

52. Conesa A, Gotz S, Garcia-Gomez J, Terol J, Talon M, Robles M. Blast2GO: a universal tool for annotation, visualization and analysis in functional genomics research. Bioinformatics. 2005;21:3674–6.

53. Schiessl K, Muiño JM, Sablowski R. Arabidopsis JAGGED links floral organ patterning to tissue growth by repressing Kip-related cell cycle inhibitors. Proc Natl Acad Sci U S A. 2014;111: 2830–2835.

54. Schuster C, Gaillochet C, Lohmann JU. Arabidopsis HECATE genes function in phytohormone control during gynoecium development. Development. 2015;142:3343–50

55. Semiarti E, Ueno Y, Tsukaya H, Iwakawa H, Machida C, Machida Y. The ASYMMETRIC LEAVES2 gene of Arabidopsis thaliana regulates formation of a symmetric lamina, establishment of venation and repression of meristem-related homeobox genes in leaves. Development. 2001;128:1771–83.

56. Simonini S, Deb J, Moubayidin L, Stephenson P, Valluru M, Freire-Rios A, et al. A noncanonical auxin-sensing mechanism is required for organ morphogenesis in Arabidopsis. Genes Dev. 2016;30:2286–2296.

57. Zumajo-Cardona C, Pabón-Mora N. Evolution of the APETALA2 Gene Lineage in Seed Plants. Mol Biol Evol. 2016;33:1818–32

58. Guo M, Thomas J, Collins G, Timmermans MCP. Direct Repression of KNOX Loci by the ASYMMETRIC LEAVES1 Complex of Arabidopsis. Plant Cell Online. 2008;20:48–58.

59. Jaradat MR, Ruegger M, Bowling A, Butler H, Cutler AJ. A comprehensive transcriptome analysis of silique development and dehiscence in *Arabidopsis* and *Brassica* integrating genotypic, interspecies and developmental comparisons. GM Crops Food. 2014;5:302–20

60. Kay P, Groszmann M, Ross JJ, Parish RW, Swain SM. Modifications of a conserved regulatory network involving INDEHISCENT controls multiple aspects of reproductive tissue development in Arabidopsis. New Phytol. 201;197:73–87

61. Marsch-Martínez N, Zúñiga-Mayo VM, Herrera-Ubaldo H, Ouwerkerk PBF, Pablo-Villa J, Lozano-Sotomayor P, et al. The NTT transcription factor promotes replum development in Arabidopsis fruits. Plant J. 2014;80:69–81.

62. Pinyopich A, Ditta GS, Savidge B, Liljegren SJ, Baumann E, Wisman E, et al. Assessing the redundancy of MADS-box genes during carpel and ovule development. Nature. 2003;424:85–8.

63. Rajani S, Sundaresan V. The Arabidopsis myc/bHLH gene alcatraz enables cell separation in fruit dehiscence. Curr Biol. 2001;11:1914–22.

64. Romera-Branchat M, Ripoll JJ, Yanofsky MF, Pelaz S. The WOX13 homeobox gene promotes replum formation in the *Arabidopsis thaliana* fruit. Plant J. 2013;73:37–49

65. Busch A, Horn S, Zachgo S. Differential transcriptome analysis reveals insight into monosymmetric corolla development of the crucifer Iberis amara. BMC Plant Biol. 2014;14:285.

66. Sinha S, Raxwal VK, Joshi B, Jagannath A, Katiyar-Agarwal S, Goel S, et al. De novo transcriptome profiling of cold-stressed siliques during pod filling stages in Indian mustard (Brassica juncea L.). Front Plant Sci. 2015;6:932

67. Balanzà V, Roig-Villanova I, Di Marzo M, Masiero S, Colombo L. Seed abscission and fruit dehiscence required for seed dispersal rely on similar genetic networks. Development. 2016;143:3372–81

68. Alonso-Cantabrana H, Ripoll JJ, Ochando I, Vera A, Ferrándiz C, Martínez-Laborda A. Common regulatory networks in leaf and fruit patterning revealed by mutations in the Arabidopsis ASYMMETRIC LEAVES1 gene. Development. 2007;134:2663–71.

69. Barker MS, Vogel H, Schranz ME. Paleopolyploidy in the Brassicales: Analyses of the Cleome Transcriptome Elucidate the History of Genome Duplications in Arabidopsis and Other Brassicales. Genome Biol Evol. 2010;1:391–9.

70. Cardinal-McTeague WM, Sytsma KJ, Hall JC. Biogeography and diversification of Brassicales: A 103 million year tale. Mol Phylogenet Evol. 2016;99:204–24.

71. Edger PP, Hall JC, Harkess A, Tang M, Coombs J, Mohammadin S, et al. Brassicales phylogeny inferred from 72 plastid genes: A reanalysis of the phylogenetic localization of two paleopolyploid events and origin of novel chemical defenses. Am J Bot. 2018;105:463–9.

72. Brock K. Tracking the Evolutionary History of Development Genes: Implications for the Diversification of Fruits and Flowers in the Brassicaceae and Cleomaceae. M.Sc. Thesis, The University of Alberta. 2014.

